# *flashfm-ivis*: interactive visualisation for fine-mapping of multiple quantitative traits

**DOI:** 10.1101/2022.03.22.485315

**Authors:** Feng Zhou, Adam S Butterworth, Jennifer L Asimit

## Abstract

**Summary:** *flashfm-ivis* provides a suite of interactive visualisation plots to view potential causal genetic variants that underlie associations that are shared or distinct between multiple quantitative traits and compares results between single- and multi-trait fine-mapping. Unique features include network diagrams that show joint effects between variants for each trait and regional association plots that integrate fine-mapping results, all with user-controlled zoom features for an interactive exploration of potential causal variants across traits.

**Availability and Implementation:** *flashfm-ivis* is an open-source software under the MIT license. It is available as an interactive web-based tool (http://shiny.mrc-bsu.cam.ac.uk/apps/flashfm-ivis/) and as an R package format. Code and documentation are available at https://github.com/fz-cambridge/flashfm-ivis and https://zenodo.org/record/6376244#.YjnarC-l2X0. Additional features can be downloaded as standalone R libraries to encourage reuse.

## Introduction

Genome-wide association studies (GWAS) have successfully identified many genetic variants that are associated with diseases and traits (Claussnitzer et al., 2020). Identifying the causal variants that underlie genetic associations is key to translating these findings into new therapeutic targets or revealing new biological insights for diseases. Statistical fine-mapping aids this by identifying potential causal variants with the aim of reducing the number of genetic variants for follow-up in downstream functional validation experiments (Spain and Barrett, 2015; Hutchinson et al., 2020). As biologically related traits often have shared causal variants, multi-trait fine-mapping that shares information between traits can improve precision over single-trait fine-mapping of each trait independently.

There are few multi-trait fine-mapping methods that allow multiple causal variants at a single locus due to the computational complexity of many possible model combinations between traits. One approach is to restrict traits to have the same causal variants and allow for different effect sizes, as in fastPAINTOR (Kichaev et al., 2017). In contrast, flashfm (Hernandez et al., 2021) makes no such restrictions and uses a Bayesian framework that upweights joint models with shared causal variants. In extensive simulation comparisons, flashfm was shown to have higher precision than fastPAINTOR.

Bioinformatics tools are moving in the direction of dynamic interaction between GWAS data and plots (**Table_S1**), but most require some programming knowledge; none of these help to explore fine-mapping results. Non-interactive fine-mapping visualisation tools include PAINTOR-CANVIS for visualising a single set of fine-mapping results (and linkage disequilibrium (LD) structure) from PAINTOR (Kichaev et al., 2017), and echolocatoR (Schilder et al., 2021) for single-trait fine-mapping results from several methods; both require some programming knowledge.

*Flashfm-ivis* provides interactive exploration and publication-ready plots to summarise fine-mapping results from multiple traits. Linked from *flashfm-ivis*, users may use *finemap-ivis* to interact with and plot single-trait results. **Table_1** compares the key features of the tools most similar to *flashfm-ivis*.

**Table_1:** Comparison of visualisation features and tools for GWAS and fine-mapping results. A “+” indicates that the tool has the feature, possibly with a few modifications.

| Features and tools | Flashfm-ivis | echolocator | PAINTOR-CANVIS | PheGWAS | LocusZoom.js | Cgmisc | LDlink | Assocplots | IntAssoPlot |
| --- | --- | --- | --- | --- | --- | --- | --- | --- | --- |
| Input data in/from different formats/methods | + |  |  |  |  |  |  |  |  |
| Display fine-mapping results | + | + | + |  |  |  |  |  |  |
| Multi-panel comparison with linked data | + | + |  |  |  |  |  | + |  |
| Joint SNP effects (fine-mapping model PPs) | + |  |  |  |  |  |  |  |  |
| Interactive features/tools | + |  |  | + | + |  | + | + |  |
| Download outputs/plots | + | + | + | + |  |  | + |  |  |
| A standalone web or R package | + |  |  | + | + |  | + |  |  |
| Regional association plot of GWAS results | + | + | + | + | + | + | + | + | + |
| Scatter plot of SNP PPs (MPP) only | +<sup>a</sup> | + | + |  |  |  |  |  |  |
| Regional association plot with SNP PPs | + |  |  |  |  |  |  |  |  |
| LD with lead SNP | + | + |  |  | + | + | +<sup>b</sup> |  | + |
| LD heatmap and plot | + |  | + |  |  |  | +<sup>b</sup> |  | + |
| Link between GWAS and LD matrix | + |  |  |  |  |  |  |  | + |
| Integrated display of multiple GWAS | + |  |  | + |  |  |  | + | + |
| No programming knowledge required | + |  |  |  | + |  | + |  |  |
| Allow iPad or touch screen | + |  |  |  |  |  |  |  |  |
| Can be used with other R/Python packages | + | + |  | + | + | + |  |  |  |
| Reference |  | Schilder et al., 2021 | Kichaev et al., 2017 | George et al., 2020 | Boughton et al., 2021; Pruim et al., 2010 | Kierczak et al., 2015 | Machiel and Chanock, 2015 | Khramtsova and Stranger, 2017 | He et al. 2020 |

<sup>a</sup>*Flashfm-ivis* integrates the SNP PPs with the regional association plots. <sup>b</sup>LDlink only uses built-in reference panels.

Flashfm multi-trait fine-mapping uses single-trait fine-mapping results from FINEMAP (Benner et al., 2016) or JAM (Newcombe et al., 2016), and we refer to either of these methods as “fm”. For ease in interpretation, variant groups are constructed in flashfm for both fm and flashfm such that variants in the same group can be viewed as exchangeable, i.e., they are in high LD and rarely appear in the same model together. Key features of *flashfm-ivis* are shown in **Fig.1** (also listed in **Table_S2)**, with an overview here:

**Fig. 1:**
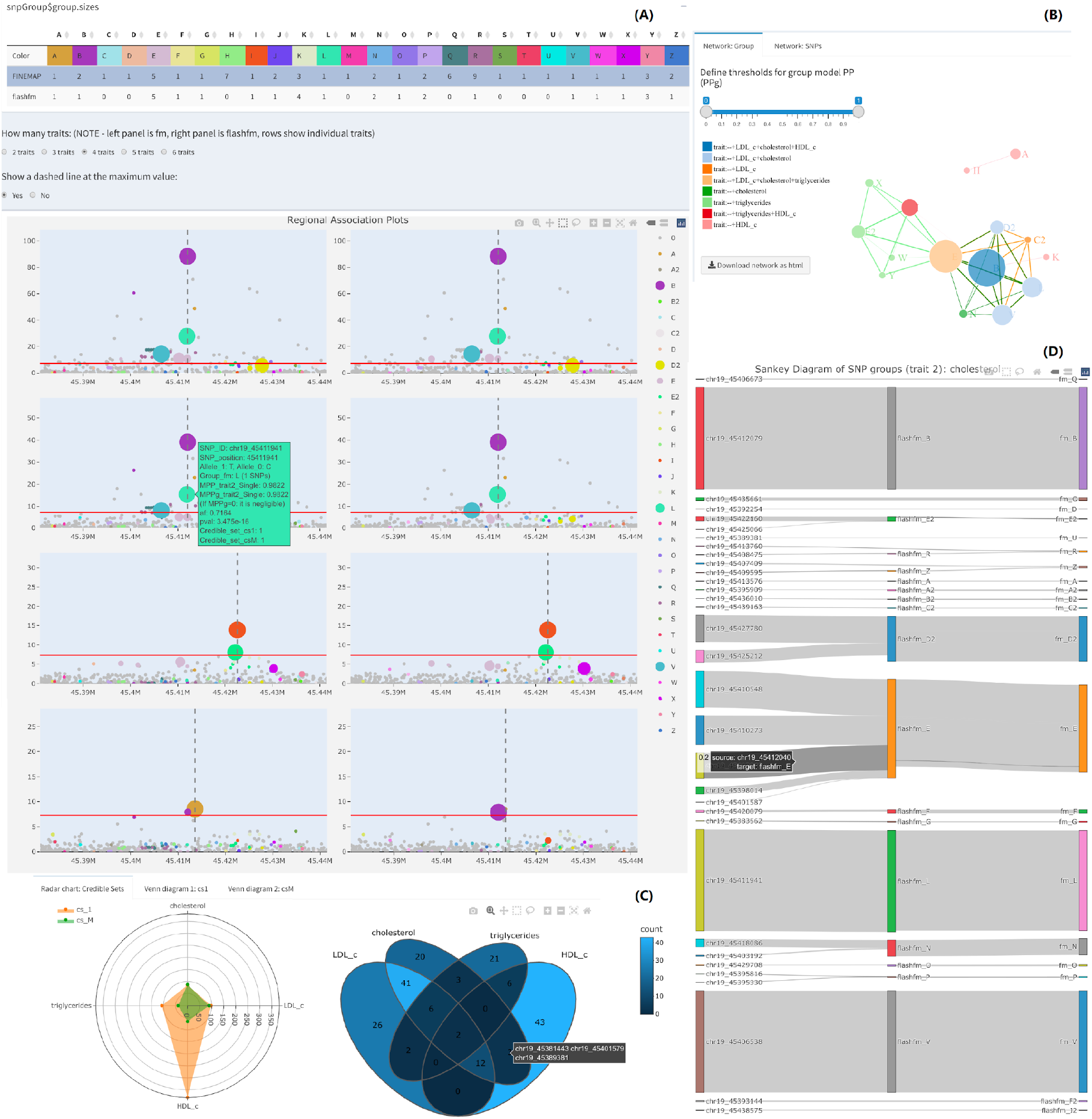
Selected *flashfm-ivis* interactive plots for 4 lipid traits from Ugandan GWAS data in the *APOE* region^a^. **(A) Interactive and Linked regional association plots with integrated fine-mapping results**. Four lipid traits (LDL-cholesterol, total cholesterol (TC), triglycerides (TG), and HDL-cholesterol) were single- (left panel) and multi-trait (right panel) fine-mapped; the rows correspond to each trait: LDL, TC, TG, HDL. The regional association plots (-log10(p) against variant position) integrate the variant MPPs (see Section 2.2) from fine-mapping, by displaying the diameter of each variant point as proportional to its MPP. Colours indicate variant groups, which consist of variants in high LD and could be treated as interchangeable in the causal models. Users may click on particular groups to focus on (or to remove) in all plots, or they may draw their selection of points in one plot as a focus for all plots. **(B) Network diagram of variant group causal models**. variant group models are displayed for the four lipid traits. Each node represents a variant group, and edges between nodes indicate variant groups that appear together in a model; only models with PPg (as defined in Section 2.4) within the range of the widget selection are included. The colours of the nodes indicate which traits include the variant group in their models, and their diameter is proportional to the frequency that they appear in models over all traits. Edge colours indicate which traits include the pair of variant groups in a model, while edge thickness is proportional to the PPg values. **(C) Radar Chart and Venn diagram of credible sets for traits**. These plots show and compare the number of variants that appear in multiple credible sets for different methods and traits. For example, in the Venn diagram the four traits share 2 variants in their credible sets constructed from flashfm, while the Radar chart compares the total number of variants based on FINEMAP (cs_1) and flashfm (cs_M). Hovering on that segment shows the variant ids, and details of each segment are in a downloadable table. **(D) Sankey diagram connects individual variant ids and their groups in flashfm or FINEMAP (fm)**. This links the individual variants and their groups in different methods. The size of the edge reflects the MPP value of the variant in each trait (e.g., trait_2 cholesterol here, users can interact with the plots to show the values). ^a^Hernandez et al. 2021.

- Interactive visualisations encourage users to compare results for multiple traits
- Downloadable summary output from different plots, based on users’ selections (e.g., credible sets, variant groups, etc.), simplifying complex results and giving insight for further analyses (e.g., discovering shared variants in credible sets for different trait combinations).
- Flexible ways of viewing the joint effects of variantS on one or more traits.

## Implementation

*Flashfm-ivis* is currently built in R and minimises the complex (inter-)dependency with other R packages (**Table_S2**), making it a standalone tool (easier to maintain in the long-term). The web-based version (http://shiny.mrc-bsu.cam.ac.uk/apps/flashfm-ivis/) does not require users to have any programming skills, encouraging a wide spectrum of researchers to interact with and visualise their own results. It includes six Dashboard tabs that give access to various comparisons and summaries, links to download all codes (https://github.com/fz-cambridge/flashfm-ivis) and a direction to *finemap-ivis* (http://shiny.mrc-bsu.cam.ac.uk/apps/finemap-ivis/).

### 2.1 Data inputs

A pre-loaded example from a cardiometabolic GWAS of a Ugandan cohort for four lipid traits and the *APOE* region (Hernandez et al. 2022) is included in *flashfm-ivis* to illustrate its main features (**Figure_S1**).

To view both single and multi-trait fine-mapping results, users need to upload their output from flashfm and its input files to *flashfm-ivis* (details in **Supplementary Material**). To view only single-trait fine-mapping results from FINEMAP (**Figure_S13)**, users upload the standard FINEMAP output files to *finemap-ivis*.

Users can verify their inputs on a sub-dashboard (**Figure_S1** and **Figure_S12**). All plots are interactive, allowing the user to control zoom-in focus areas in regional association plots and to drag network nodes to change the perspective of plots. An overview of key plots follows (also see **Table_S2**, all detailed plots in **Supplementary Material**).

### 2.2 Control widgets on the sidebar panel

Control widgets help users to refine the input data selection and then focus on key results; both credible sets and *MPP* (marginal posterior probability of variant causality – the posterior probability that the variant is included in a model) values can be controlled (**Figure_S2**).

### 2.3 Fine-map integrated regional association plots

Individual regional association plots are presented for each trait in **Figure_S2** and MPPs from fine-mapping are denoted by the dot size. In **Figure_S8**, all regional association plots are linked together for easy comparisons between traits and between methods – users may select a subset of variants to focus on in all plots.

### 2.4 Network plots of joint variant effects

**Figures S4-S7** present both variant group- and individual-level dynamic networks. The key features are: (1) users interact with the networks by specifically defining the PP thresholds for variant models, so the network can be expanded or refined; (2) the size and colour of both nodes and edges indicate evidence strength and associated traits, as detailed in **Table_S2**; (3) sub-networks can be formed and moved. When viewing models based on variant groups, we consider the group model PP (PPg), which is the sum of the PPs for all models that have exactly one variant from each group listed in the model; e.g., model A+B consists of all variant models with one variant from each of groups A and B (i.e. fm in “Ch_1_Single_trait’’ and flashfm in “Ch_2_Multi_trait”).

### 2.5 Venn diagrams of shared potential causal variants

In **Fig.1** and **Figure_S10**, through interactive Venn diagrams, users see and download lists of variants in each combination of overlapping credible sets.

### 2.6 Spider/radar diagram of credible set sizes

In **Fig.1** and **Figure_S9**, the spider chart compares the number of variants in credible sets of fm/flashfm methods based on different traits.

### 2.7 Sankey diagrams of variant groups

In **Fig.1** and **Figure_S11**, Sankey flowcharts show the connections between variants and variant groups in fm/flashfm methods and based on different traits.

## Conclusion

We provide a user-friendly fully interactive web tool, *flashfm-ivis*, that is accessible without any programming knowledge; it is available as an R package for those who prefer to run it from their own machine. It promotes exploration of fine-mapping results of several traits and helps with interpretation, as well as identification of variants for further follow-up. Users can interact with plots and decide on the final version of publication-ready plots for download. *Flashfm-ivis* output, such as lists of variants in credible sets for selected traits, is easily downloaded for further follow-up. If users are interested in only a single trait, there is an option to produce plots based on FINEMAP output only, *finemap-ivis*. We hope that *flashfm-ivis* will become standard practice for exploring fine-mapping results and will contribute to revealing the underlying mechanisms of diseases.

## Acknowledgements

The authors thank Jana Soenksen and Inês Barroso, both from the Exeter Centre of Excellence for Diabetes Research (EXCEED), University of Exeter Medical School, Exeter, UK, for their valuable feedback and suggestions.

## Data Availability

The example datasets can be found in the web-based version of flashfm.ivis (http://shiny.mrc-bsu.cam.ac.uk/apps/flashfm-ivis/) and its GitHub repository (https://github.com/fz-cambridge/flashfm-ivis), as well as https://zenodo.org/record/6376244#.YjnarC-l2X0.

## Supplementary Material

See the online appendix and the README tab of *flashfm-ivis* (http://shiny.mrc-bsu.cam.ac.uk/apps/flashfm-ivis/), which also includes YouTube video demonstrations of the tool.

