## Supplementary Material for "*flashfm-ivis*: interactive visualisation for fine-mapping of multiple quantitative traits"

**SUPPLEMENTARY TABLE_S1.** Other available GWAS visualisation software packages

| Name of tools | Software link | Citation | Purpose and description |
|---|---|---|---|
| Assocplots | https://github.com/khramts/assocplots | Khramtsova and Stranger, 2017, *Bioinformatics* | A python package for static and interactive visualisation of multiple-group GWAS results |
| cgmisc | https://github.com/cgmisc-team/cgmisc | Kierczak et al., 2015, *Bioinformatics* | An R package that enables enhanced data analysis and visualisation of results from GWAS |
| echolocatoR | https://github.com/RajLabMSSM/echolocatoR | Schilder et al., 2021, *Bioinformatics* | Automated statistical and functional fine-mapping with extensive access to genome-wide datasets |
| FIVEx | https://fivex.sph.umich.edu<br>https://github.com/statgen/fivex/ | Kwong et al., 2021, *Bioinformatics* | A web browser tool for visualising Genotypes, expression and splice QTL (cis-eQTL and cis-sQTL) datasets |
| GAPIT | https://www.maizegenetics.net/gapit<br>https://github.com/jiabowang/GAPIT3 | Lipka et al., 2012, *Bioinformatics*<br>Wang and Zhang, 2021, *Genomics, Proteomics and Bioinformatics (GPB)* | An R package that performs a GWAS and genome prediction (or selection) |
| GSDS | http://gsds.cbi.pku.edu.cn | Hu et al., 2015, *Bioinformatics* | Visualizing genes' structure and annotated features helps biologists to investigate their function and evolution intuitively |
| Haploview | https://www.broadinstitute.org/haploview/haploview | Barrett et al., 2005, *Bioinformatics* | Haploview is designed to simplify and speed up |

| | | | |
|---|---|---|---|
| | | | haplotype analysis by providing a common interface to several tasks relating to such analyses |
| IntAssoPlot | https://github.com/whweve/IntAssoPlot | He et al., 2020, *frontiers in Genetics* | Visualize GWAS with Gene Annotation and Linkage Disequilibrium |
| JBrowse | https://jbrowse.org/jb2/ | Buels et al., 2016, *Genome Biology* | A fast and full-featured genome browser built with JavaScript and HTML5 |
| LDlink | https://ldlink.nci.nih.gov/?tab=home https://github.com/CBIIT/nci-webtools-dceg-linkage | Machiela and Chanock, 2015, *Bioinformatics* | A suite of web-based applications designed to interrogate LD easily and efficiently in population groups |
| LocusZoom | http://locuszoom.org | Pruim et al., 2010, *Bioinformatics* | A suite of tools to provide fast visualization of GWAS results for research and publication |
| LocusZoom.js | https://github.com/statgen/locuszoom/releases/tag/v0.12.0 | Boughton et al., 2021, *Bioinformatics* | A JavaScript implementation of LocusZoom to make LocusZoom embeddable and interactive |
| PAINTOR-CANVIS | https://github.com/gkichaev/PAINTOR_V3.0/tree/master/CANVIS | Kichaev et al., 2017, *Bioinformatics* | A visualisation tool utilized for the output of PAINTOR |
| PheGWAS | https://github.com/georgeg0/PheGWAS | George et al., 2020, *Bioinformatics* | Three-dimensional approach to dynamically visualize GWAS across multiple phenotypes |
| ShinyGPA | https://github.com/dongjunchung/GPA | Kortemeier et al., 2018, *PLoS One* | An interactive visualization toolkit for investigating pleiotropic architecture using GWAS datasets |
| Zbrowse | https://github.com/baxterlabZbrowse/ZBrowse | Ziegler et al., 2015, *PeerJ Comput. Sci.* | An interactive GWAS viewer focused on comparing results |

| | | | |
|---|---|---|---|
| | | | across traits and other variables |

**SUPPLEMENTARY TABLE_S2.** Summary of key functions implemented in *flashfm-ivis*

| Dashboard | Charts | Functions | Features |
|---|---|---|---|
| GWAS summary statistics (Ch_0 Input Values)<br>**See Figure S2 and S3** | Interactive regional association plots | + Colour: SNP Group<br>+ Size: MPP value<br>+ X axis: SNP location<br>+ Y axis: P-value<br>+ Widget_1: credible sets selection<br>+ Widget_2: define MPP value<br>+ Legend: user selection of SNP Group | Interactive tools:<br>+ Download plots directly<br>+ Box or Lasso Selection<br>+ Zoom In and Out<br>+ Show and compare data on hover<br>+ Data selection via legend element<br>+ Links to extra widgets |
| | Coloured SNP Group | + Colour: link to the SNP Group in plots<br>+ Value: number of SNPs in group | Possible implementation: user can download the table directly |
| | Interactive and selective LD plot | + Heatmap: an initial overview of LD<br>+ LD plot: show selected SNPs | Interactive and dynamic features |
| Single trait fine mapping network (Ch_1 Single trait)<br>**See Figure S4 and S5** | Group-based network<br>Individual SNP-based network | + Widget: define PP values<br>+ Colour of node: trait combination sets<br>+ Size of node: frequency in each set<br>+ Thickness of edge: PP values<br>+ Colour of edge: sub-network of trait | Interactive tools:<br>+ Define PP values<br>+ Understanding sub-networks based on colours, nodes and links |
| Multi trait fine mapping network (Ch_2 Multi trait)<br>**See Figure S6 and S7** | Group-based network<br>Individual SNP-based network | | |
| Linked data comparison plots (Ch_3 Comparison (Linked))<br>**See Figure S8** | Coloured SNP Group<br>Coloured and linked regional association plots | + Compare linked regional association plots across all different traits<br>+ All colours of SNP Group are consistent in different plots<br>+ Users' own selections of data | Combined all interactive tools as above, but also linking selected data of different plots together |

| | | | |
|---|---|---|---|
| Method comparison plots (Ch_4 Comparison (Methods))<br><br>**See Figure S9 and S10** | Radar chart of credible sets<br><br>Venn diagram of credible sets | + Comparison of credible sets from different methods<br>+ Show individual SNPs in different joint sets | Allow users to interact with the diagrams and understand (as well as download) individual SNPs in each joint set |
| Trait comparison plots (Ch_5 Comparison (Traits))<br><br>**See Figure S11** | All trait combined Sankey diagram<br><br>Individual traits of Sankey diagram | + Comparison of SNP groups in different traits<br>+ Show detailed connections between different groups in two methods | Allow users to focus on the main SNPs in particular SNP Groups based on different methods |

NOTE: This initial version of flashfm-ivis is built completely in R but can be extended to other platforms (such as Python) for a more diversified audience with different backgrounds. It is designed to minimise the complex (inter-)dependency with other R packages, making it a standalone tool, so it will be easier to maintain in the long-term. Some functions require a few existing commonly-used open-source packages in the R community (see CRAN.R-project.org): plotly (Sievert, 2020; https://plotly.com/r/), ggplot2 (Wickham, 2011; https://ggplot2.tidyverse.org), shiny (Verity, et al., 2017; https://shiny.rstudio.com and https://cran.r-project.org/package=shiny), shinydashboard (Chang, 2015; https://rstudio.github.io/shinydashboard/), DT (https://rstudio.github.io/DT/), gaston (https://cran.r-project.org/package=gaston), randomcoloR (https://cran.r-project.org/package=randomcoloR), networkD3 (Allaire, et al., 2017; https://cran.r-project.org/package=networkD3), ggVennDiagram (Gao, et al., 2021; https://cran.r-project.org/package=ggVennDiagram), dplyr (https://dplyr.tidyverse.org), igraph (Csardi and Nepusz, 2016; https://igraph.org/r/), stringr (https://cran.r-project.org/package=stringr).

**Flashfm-ivis (FLexible And Shared information Fine-Mapping-Interactive VISualisation)**

### *Instructions: Input files*

The key variables and their names should be fixed as follows:

Finemap-ivis: http://shiny.mrc-bsu.cam.ac.uk/apps/finemap-ivis/
Users can upload the four standard input/output files from FINEMAP (Benner et al. 2016). They can use any names for the files, but the file extensions should be fixed as follows:

1. Anyname.config
2. Anyname.ld
3. Anyname.snp
4. Anyname.z

Flashfm-ivis: http://shiny.mrc-bsu.cam.ac.uk/apps/flashfm-ivis/
This requires output from running multi-trait fine-mapping with flashfm (Hernandez et al. 2021).
Depending on the following three situations (i.e. options on the webpage sidebar), users can upload different data files by using "Browse" button:

1. **Single RData file:** if users save all variables (i.e. items 1.a – 1.d) below in a single RData file, they can upload this file directly to the server, but please make sure the list and variable names are fixed as follows:
    a. A list named GWAS, where GWAS[[m]] is the data.frame for m$^{th}$ trait. Inside each data.frame GWAS[[m]], the variables (columns) must be given in order, they are:
        1) $rs: name or rsid of SNP
        2) $chr: number of chromosome (to mark the x-axis label in the plots)
        3) $ps: base-pair position of SNP
        4) $allele1: allele1 of SNP (effect allele)
        5) $allele0: allele0 or allele2 of SNP (non-effect allele)
        6) $beta
        7) $se
        8) $pval
        9) $af: effect allele frequency or minor allele frequency (MAF)
        10) $no: sample size at each SNP (i.e. if unavailable, assign zero or NA to this column)
    b. A matrix (i.e. not a list) of LD with the SNP names in colnames and rownames, matching the GWAS[[m]]$rs above.
    c. A list of flashfm output mpp.pp, which contains the 4 usual sub-lists as:
        1) mpp.pp$MPP
        2) mpp.pp$MPPg
        3) mpp.pp$PP
        4) mpp.pp$PPg
    d. A list of flashfm output snpGroups, which contains the 3 usual sub-lists as:
        1) snpGroups$groups.fm

        2) snpGroups$groups.flashfm
        3) snpGroups$group.sizes
    e. PLEASE NOTE: users can use save(GWAS, LD, mpp.pp, snpGroups, file="filename.RData") to keep/store the above four datasets together from their RStudio to one single .RData file and save it in their local working directory, then upload it to the webpage server.
    f. YouTube video for an illustration will be added on the web-based version.
2. **FINEMAP with flashfm:** if users have separate files from different sources (e.g. FINEMAP and flashfm packages), then the files with different extensions must have the following variables with the fixed names:
    a. For each trait, a standard .z file from FINEMAP for all individual traits, (please do not change the variable names, but the order of columns can be changed), which contains:
        1) $rsid
        2) $chromosome
        3) $position
        4) $allele1
        5) $allele2
        6) $maf
        7) $beta
        8) $se
    b. One .ld file that contains the standard LD matrix from FINEMAP (please do not change anything).
    c. One .RData file that was created and saved from the flashfm R package, which contains the mpp.pp and snpGroups datasets. (Please save both datasets directly in the .RData file by using save(mpp.pp, snpGroups, file="filename.RData"), do not combine them as a list).
    d. PLEASE NOTE: if users upload multiple .z files at the same time, please order the files from trait_1 to trait_m (regardless of the names of these .z files), since R-Shiny will read the first .z file sending to the server as trait 1, the second .z file as trait 2 and so on. (See YouTube video for a detailed illustration.)
    e. YouTube video for an illustration will be on the web-based version.
3. **FLASHFMwithJAM:** similar to **Finemap with flashfm,** but use the expanded JAM method in flashfm R package and create two different mpp.pp and snpGroups datasets when saving them to the .RData file). A similar YouTube video will be added on the web-based version.
4. **Single RData or csv file:** If users have a single file (i.e., .rdata or .csv formats) contains all lists and variables as mentioned above.

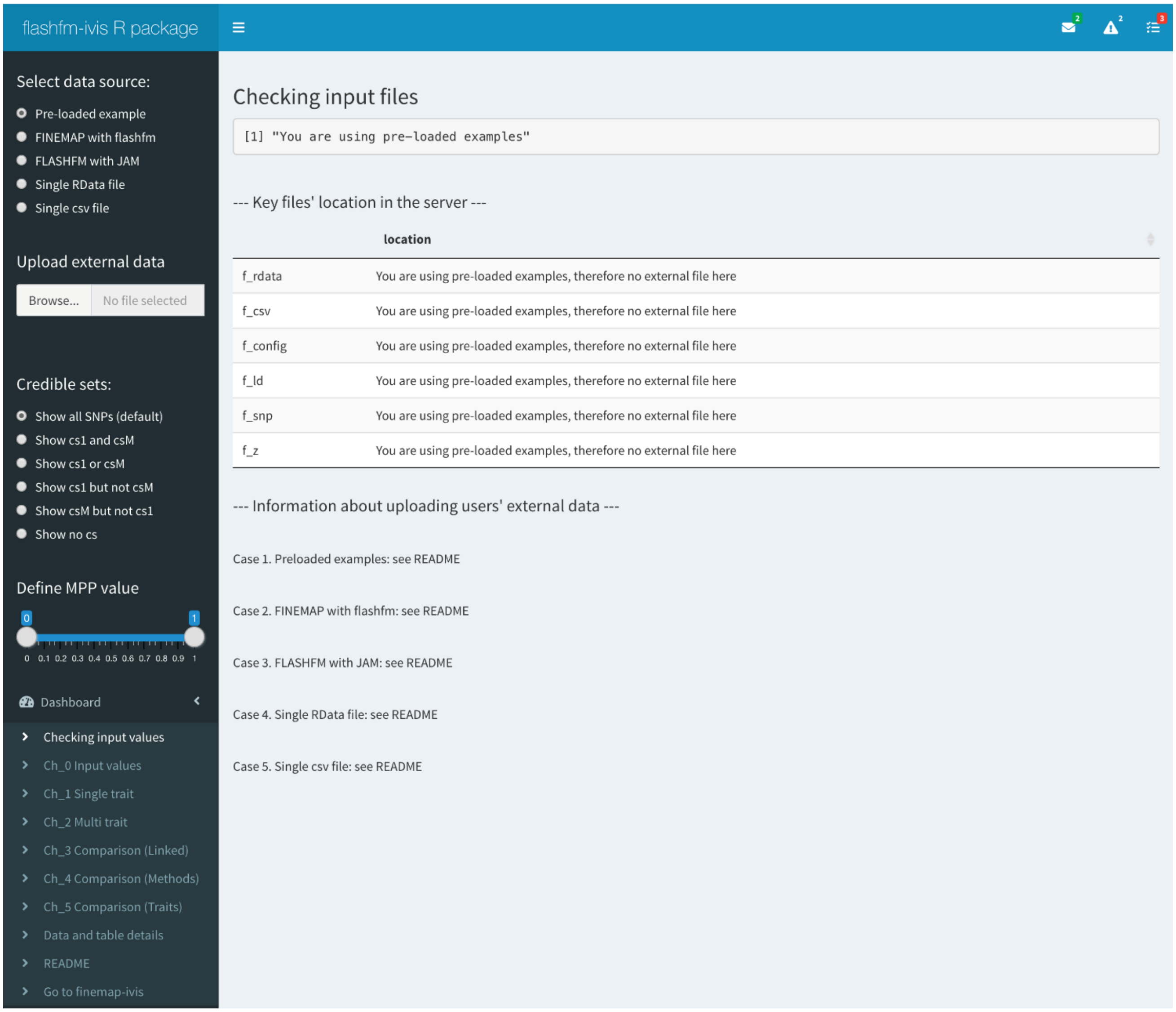

*Figure_S1: Data input page*

#### *Instructions: Available plots and data summaries*

All plots that are generated could be downloaded directly in a publication-ready format (the default type is a PNG file of size 700 x 450 pixels). The file specifics may be controlled via the toImageButtonOptions configuration key. Possible formats: 'format': 'svg', # one of png, svg, jpeg, webp. More information: [https://plotly.com/python/configuration-options/#customizing-modebar-download-plot-button](https://plotly.com/python/configuration-options/#customizing-modebar-download-plot-button).

The user may also interact with the plots to select different perspectives and download their selected version.

Below we refer to:

- **PP** (posterior probability) for multi-SNP models
- **MPP** (marginal posterior probability) is the PP for a SNP's inclusion in a causal model; it is the sum over all model PPs that include the SNP.
- **SNP groups:** these are output from the flashfm R package and are constructed from LD matrix and fine-mapping results; SNPs belonging to the same group can be viewed as exchangeable. SNPs with MPP>0.001 are assigned to the same group if they have high LD (pairwise $r^2$>0.6) and rarely appear in a model together. Groups based on flashfm tend to be subsets of those from single-trait fine-mapping.
- **PPg (PP based on SNP groups):** for a particular SNP group model A+B, PPg is the sum of PP for all models composed of exactly one SNP from group A and one SNP from group B.
- **MPPg (MPP based on SNP groups):** MPPg for group A is the sum of MPP for all SNPs that belong to group A.
- **fm** refers to a single-trait fine-mapping method JAM (Newcombe et al. 2016) if using internal function FLASHFMwithJAM or FINEMAP (http://www.christianbenner.com/ ) if using option 3 of running flashfm (https://jennasimit.github.io/flashfm/ )

**Modebar icons that offer users different interactive features in dashboard plots**

| Icon | Use |
|---|---|
| 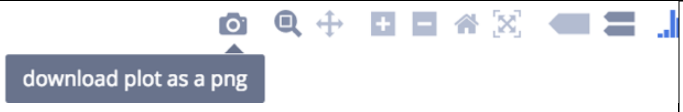  | Click on the camera icon and get/download the plot in PNG format. |
| 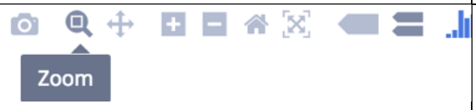  | Clicking this icon selects the Zoom mode. To zoom in on a region of a graph, click and hold your mouse, moving across the region. Release your mouse. To return to the original view, double-click anywhere on the plot. |
| 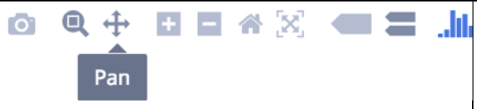  | To pan across regions of your graph, select the Pan mode. Click and hold your mouse to explore the data. Double-click anywhere to return to the original view. |
| 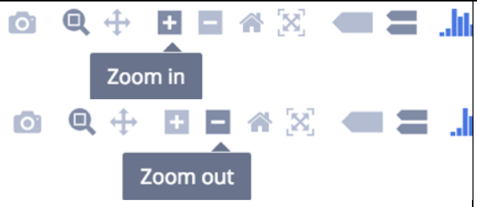  | You can zoom in and out by clicking on the + and - buttons. The plot keeps axes labels and annotations the same size to preserve readability. |
| 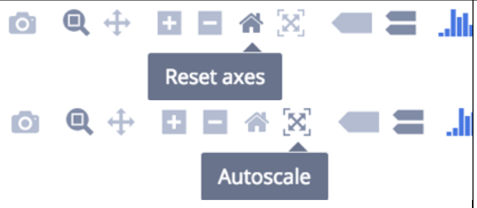  | Clicking this icon zooms to include your Axes Range, if this has been set. If it has not been set it zooms to a setting that is optimized to include all the viewable data, the same as if Autoscale had been clicked. |
| 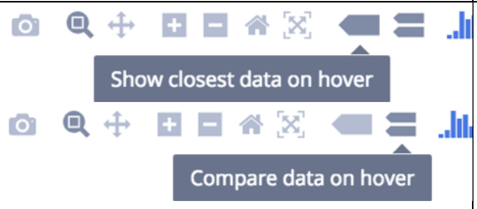  | One of these two buttons is selected at all times. Clicking 'Show closest data on hover' will display the data for just the one point under the cursor. Clicking 'Compare data on hover' will show you the data for all points with the same x-value. |

| | |
|---|---|
| 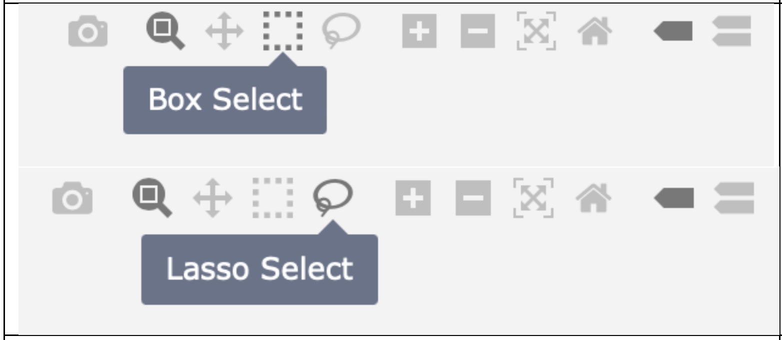 | They are basic selection tools. Box Select is to select data on the plot by creating a box, while the Lasso Select is used to draw freehand selection. |
| | **In general, by double-clicking anywhere on the plot, users can return to the original view.** |
| **More reference:** | [https://plotly.com/chart-studio-help/getting-to-know-the-plotly-modebar/](https://plotly.com/chart-studio-help/getting-to-know-the-plotly-modebar/) |

**Ch_0 Input Values – GWAS summary statistics and input values**

- **Interactive regional association plots with integrated fine-mapping results – easily view SNPs based on both genetic association and probability of causality (from single-trait and multi-trait fine-mapping) in a familiar GWAS format**

The upper panel is for single-trait fine-mapping results and the lower panel is for multi-trait fine-mapping results. Within each panel, there is a tab for each trait. Within each panel tab a regional assocation plot (-$\log_{10}$(p) against SNP position) is displayed for each trait with the following additional features:

a) Hover over a point to see SNP details (SNP ID, alleles, allele frequency, etc. Please note, if MPPg is negligible, we show it is equal to 0.)
b) Colour of points: SNP group membership according to fine-mapping results. SNPs belonging to the same group can be viewed as exchangeable. SNPs with MPP>0.001 are assigned to the same group if they have high LD (pairwise $r^2$>0.6) and rarely appear in a model together.
c) Size of points: proportional to fine-mapping posterior probabilities of SNP causality (referred to as MPP – Marginal Posterior Probability)
d) 99% credible set selection widget (see left sidebar 'Credible sets') – highlight SNPs based on inclusion/exclusion in credible sets from single-trait (i.e. defined as 'cs1' on the left sidebar) and/or multi-trait fine-mapping (i.e. defined as 'csM' on the left sidebar); can be combined with MPP widget
e) MPP range widget (see left sidebar 'Define MPP value') – highlight SNPs that have MPP falling between the minimum and maximum sliders on the widget; can be combined with credible set widget
f) Click on "Box Select" or "Lasso Select" to draw a box or a lasso (free drawing of any shape) around points to focus on and fade other points.
g) Click on "Zoom" (or Zoom In/out), then draw a box around points to zoom in and out for point selection.
h) Single click on SNP group in legend to remove all points belonging to that group.
i) Double click on SNP group in legend to remove all points not belonging to that group.
j) Click on "Pan" and then drag plot to left or right to change centre of plot.
k) Click on reset axis to res-set to default plot
l) "Download plot as a PNG" option can download the plot to the local machine
m) "Autoscale" to adjust the view of the current plot

- **Coloured SNP Groups – colours match those in the legend of the interactive regional association plots**

a) view SNP group sizes from both single and multi-trait fine-mapping
b) The table will be adjusted automatically depending on the computer screen size, but users can also use mouse/touchpad to scroll left or right of the table.

- **Interactive and selective LD plot – heat map style plot of $r^2$ between SNPs**

a) User selects which SNPs to include in LD plot based on the credible set and MPP widgets that control the interactive regional association plots (see c and d of interactive regional association plots)
b) Click on Zoom to focus on an area of the plot.

c) Other interactive features (e.g. Pan, Autoscale, Download, etc) are same as in the regional association plots.

- **Traditional LD.plot if the selected SNP sample is small, but it will show a histogram if the SNP sample is large (since the LD.plot function takes a long time to run the plot with a large sample)**
  a) Users can download the plot by clicking the button
  b) The plot will be updated according to the control options in the sidebar

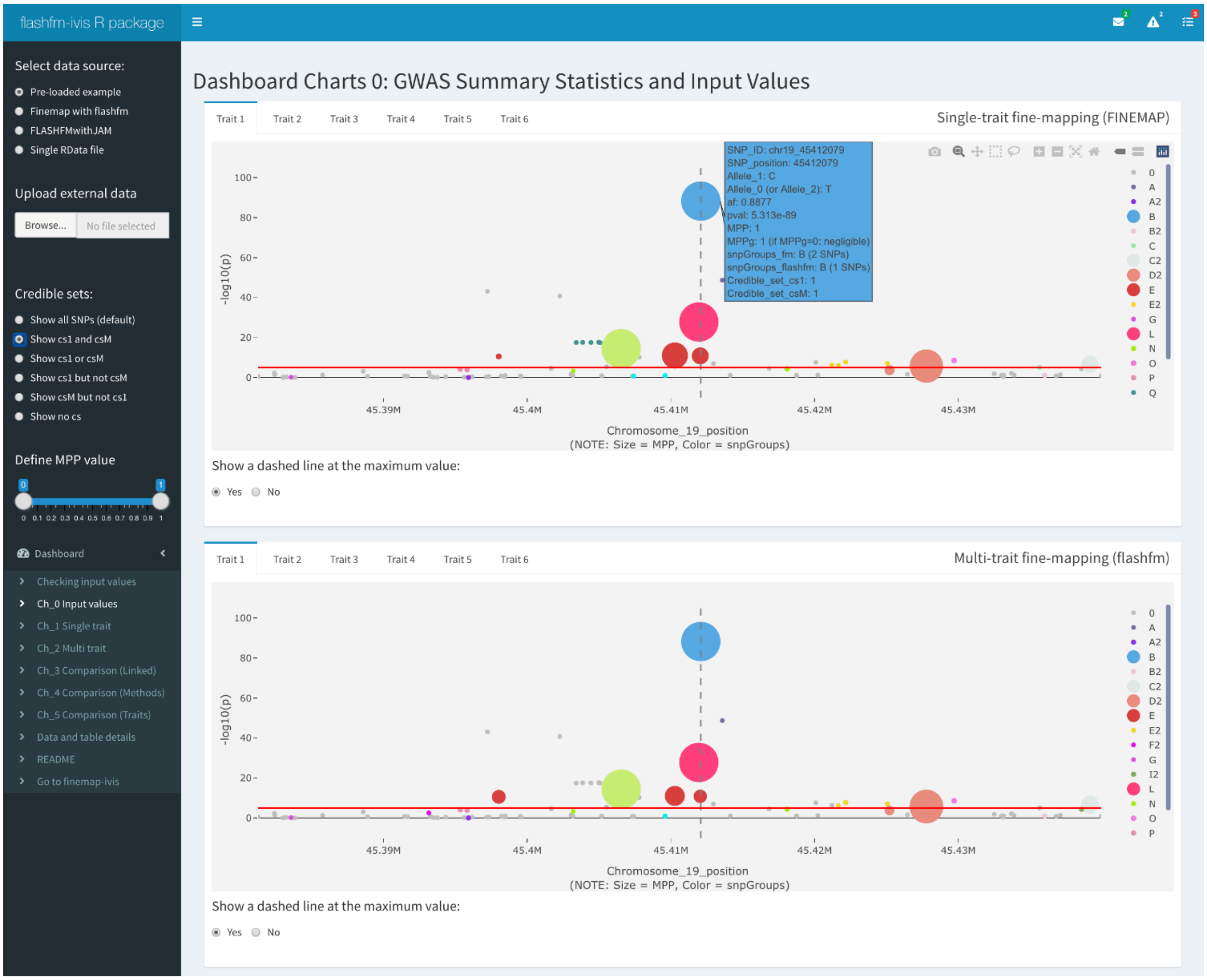

*Figure_S2: Ch_0 Input values (a)*

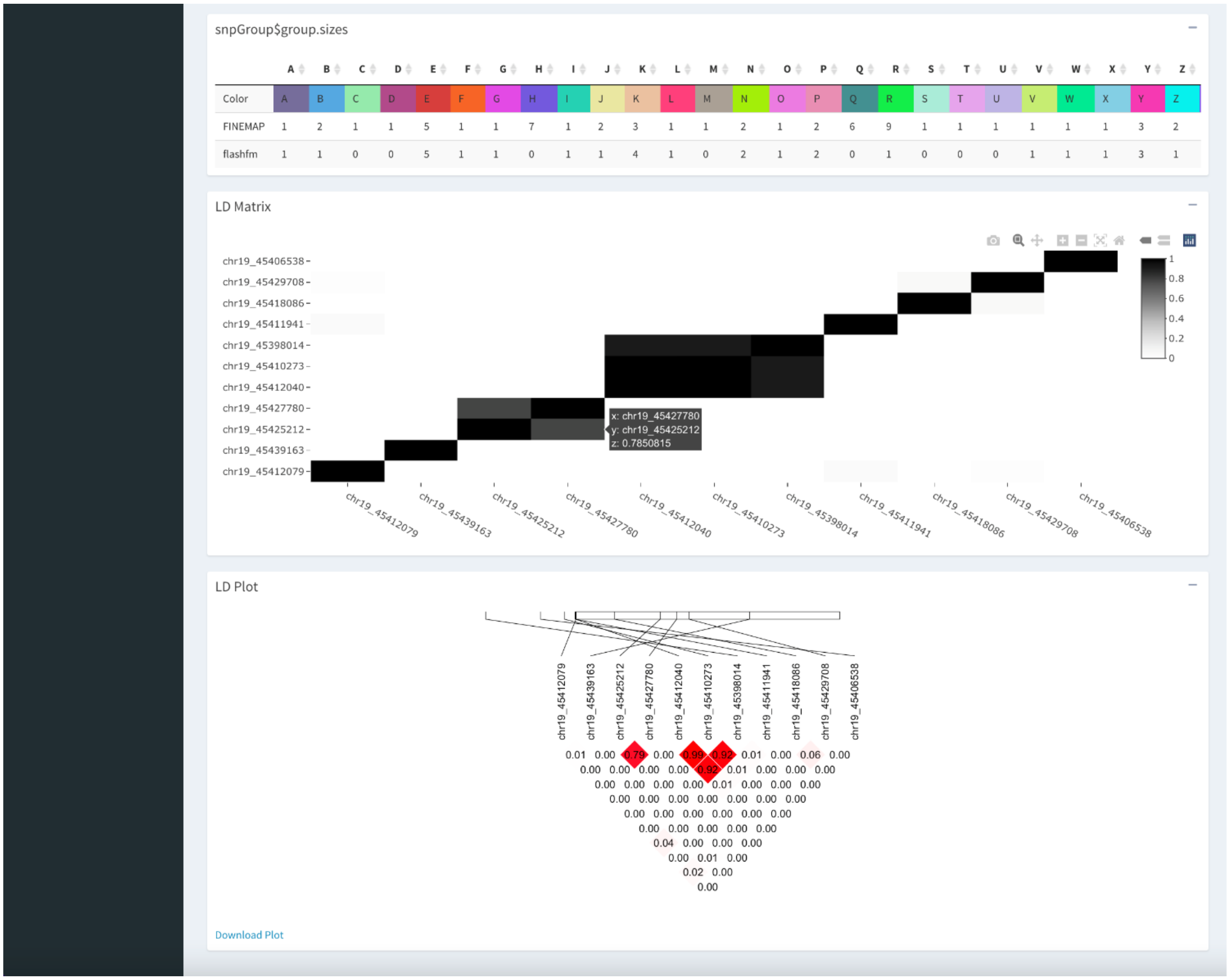


| | A | B | C | D | E | F | G | H | I | J | K | L | M | N | O | P | Q | R | S | T | U | V | W | X | Y | Z |
|---|---|---|---|---|---|---|---|---|---|---|---|---|---|---|---|---|---|---|---|---|---|---|---|---|---|---|
| Color | A | B | C | D | E | F | G | H | I | J | K | L | M | N | O | P | Q | R | S | T | U | V | W | X | Y | Z |
| FINEMAP | 1 | 2 | 1 | 1 | 5 | 1 | 1 | 7 | 1 | 2 | 3 | 1 | 1 | 2 | 1 | 2 | 6 | 9 | 1 | 1 | 1 | 1 | 1 | 1 | 3 | 2 |
| flashfm | 1 | 1 | 0 | 0 | 5 | 1 | 1 | 0 | 1 | 1 | 4 | 1 | 0 | 2 | 1 | 2 | 0 | 1 | 0 | 0 | 0 | 1 | 1 | 1 | 3 | 1 |

*Figure_S3: Ch_0 Input values (b)*

**Ch_1 Single trait – Single-trait fine-mapping models**

**Group-based network – interactive visualisation of single-trait fine-mapping PP from models, to see which SNP groups tend to appear together in a model, having joint effects on a trait(s)**

a) Colour of node shows which traits include the SNP group in their models
b) Size of node is proportional to the frequency that the SNP group appears in models
c) Thickness of edges joining nodes are proportional to the PPg values of models that include joint effects of the two SNP group nodes
d) Colour of edges indicate the sub-network of the trait, i.e., which traits have models that include the two SNP group nodes joined by the edge
e) Widget controls the range of PPg for models to display in the network
f) Nodes may be dragged to change the perspective of the plot
g) Users can scroll (Zoom in/out) the view of networks (i.e., to make the view larger or smaller to see the whole picture). Also, the view can be adjusted automatically depending on the size of the computer/tablet screen.
h) Users can download the network by clicking the button. Due to the interactive feature of this network, the downloaded plot is a dynamic html format/webpage, but users can open this html file in their local machines and save/print the network as a static PDF file or use screenshot to save it as a static PNG file.

**Individual SNP-based network – interactive visualisation of single-trait fine-mapping PP from models, to see which SNPs tend to appear together in a model, having joint effects on a trait(s)**

a) Same features as for the SNP group network
b) As there are many more SNP nodes than SNP group nodes, it is advised to have a higher minimum PP to simplify and focus on the most likely models
c) There may be more than one sub-group network (e.g., depending on the PP value, two or more separate networks may be formed), therefore users can use their mouse or touchpad to scroll (Zoom in/out) of the view, in order to see all sub-networks.
d) Since it is a large network, SNPs that are connected in a network (depending on the selected PP value) will be placed further from the centre, to focus on the main network.
e) Users can download the network by clicking the button. Due to the interactive feature of this network, the downloaded plot is a dynamic html format/webpage, but users can open this html file in their local machines and save/print the network as a static PDF file or use screenshot to save it as a static PNG file.

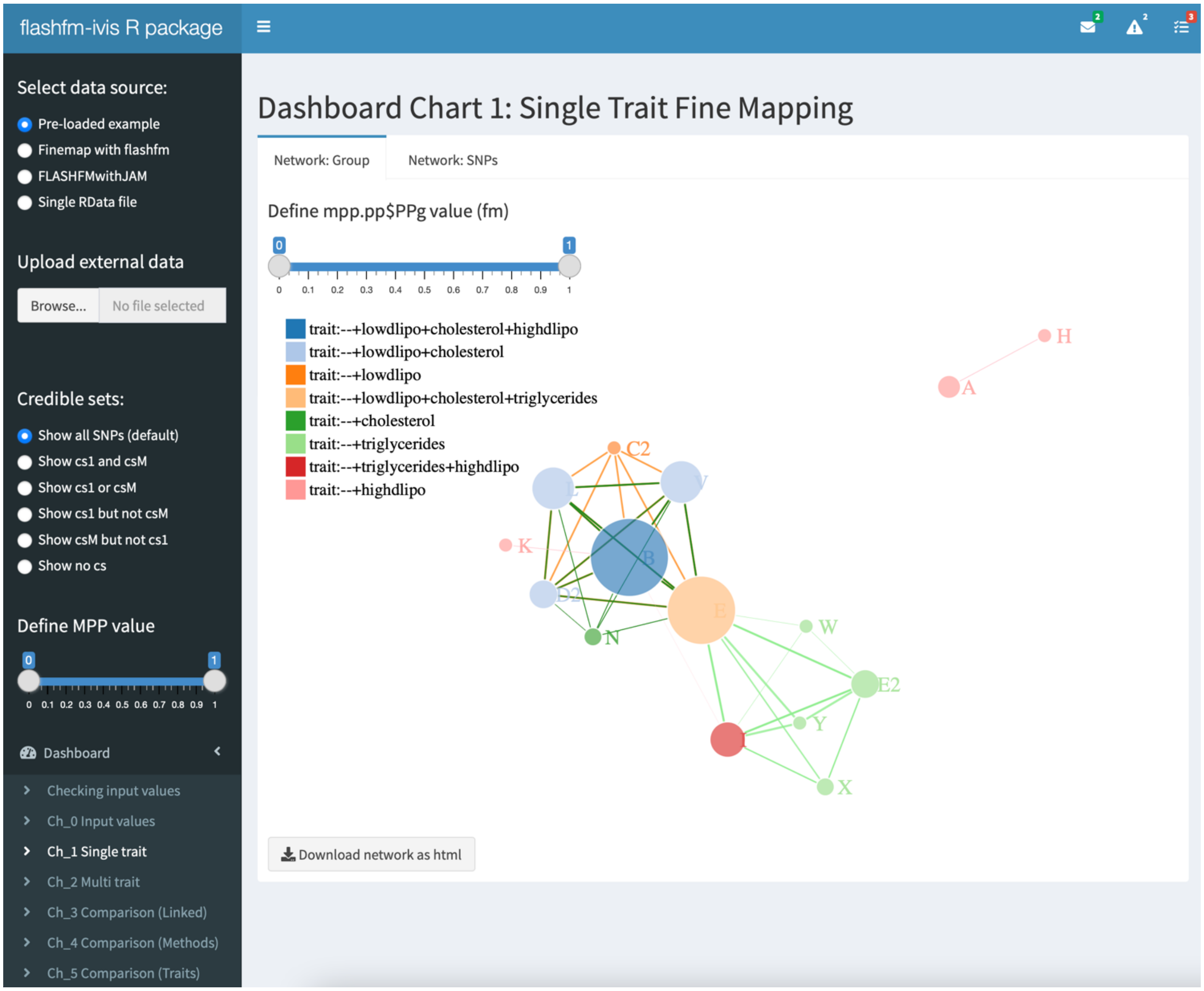

Figure_S4: Ch_1_Single_trait (a)

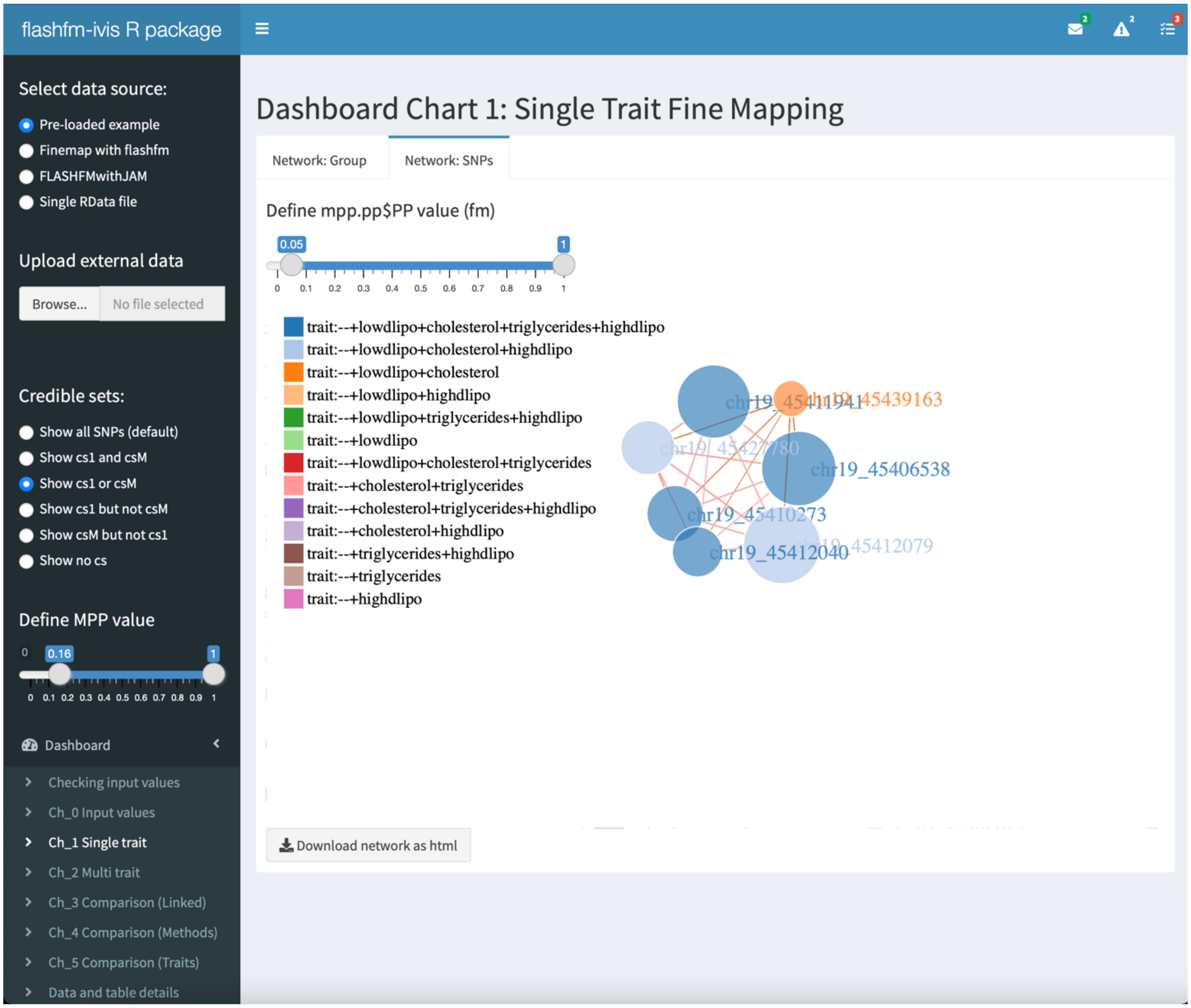

Figure_S5: Ch_1_Single_trait (b)

**Ch_2 Multi trait – Multi-trait fine-mapping models**

**Group-based network- interactive visualisation of multi-trait fine-mapping PP from models, to see which SNP groups tend to appear together in a model, having joint effects on a trait(s)**

a) Colour of node shows which traits include the SNP group in their models
b) Size of node is proportional to the frequency that the SNP group appears in models
c) Thickness of edges joining nodes are proportional to the PPg values
d) Colour of edges indicate the sub-network of the trait, i.e., which traits have models that include the two SNP group nodes joined by the edge
e) Widget controls the range of PPg for models to display in the network
f) Nodes may be dragged to change the perspective of the plot
g) Users can use their mouse or touchpad to scroll (Zoom in/out) the view of networks (i.e., to make the view larger or smaller to see the whole picture). Also, the view can be adjusted automatically depending on the size of the computer/tablet screen.
h) Users can download the network by clicking the button. Due to the interactive feature of this network, the downloaded plot is a dynamic html format/webpage, but users can open this html file in their local machines and save/print the network as a static PDF file or use screenshot to save it as a static PNG file.

**Individual SNP-based network- interactive visualisation of multi-trait fine-mapping PP from models, to see which SNPs tend to appear together in a model, having joint effects on a trait(s)**

a) All the same features as for the SNP group network
b) As there are many more SNP nodes than SNP group nodes, it is advised to have a higher minimum PP to simplify and focus on the most likely models
c) There may be more than one sub-group network (e.g., depending on the PP value, two or more separate networks may be formed), therefore users can use their mouse or touchpad to scroll (Zoom in/out) of the view, in order to see all sub-networks.
d) Since it is a large network, SNPs will move far away from the centre if they are not connected in a network (depending on the selected PP value), in order to make clear about the main network.
e) Users can download the network by clicking the button. Due to the interactive feature of this network, the downloaded plot is a dynamic html format/webpage, but users can open this html file in their local machines and save/print the network as a static PDF file or use screenshot to save it as a static PNG file.

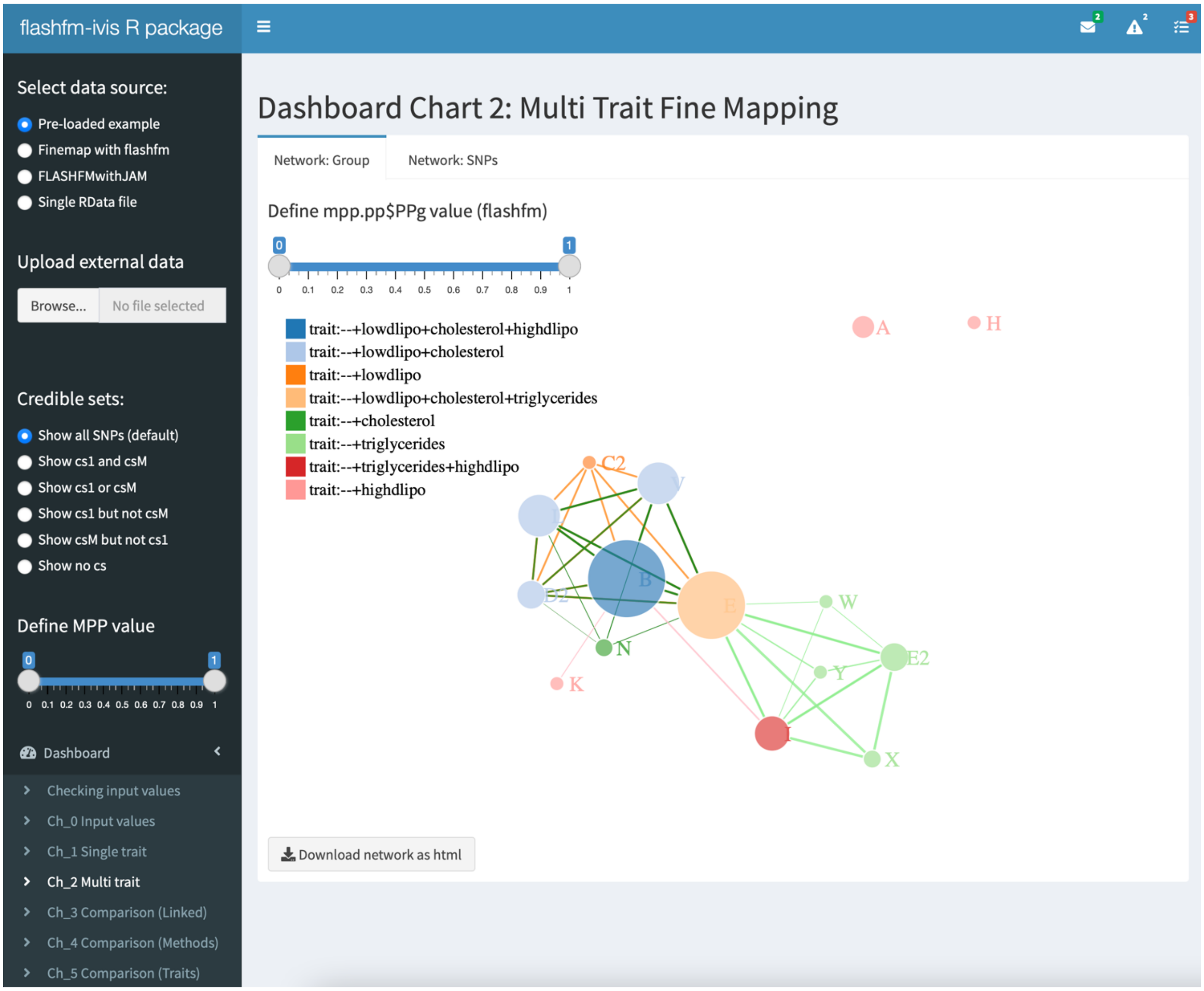


*Figure_S6: Ch_2_Multi_trait (a)*

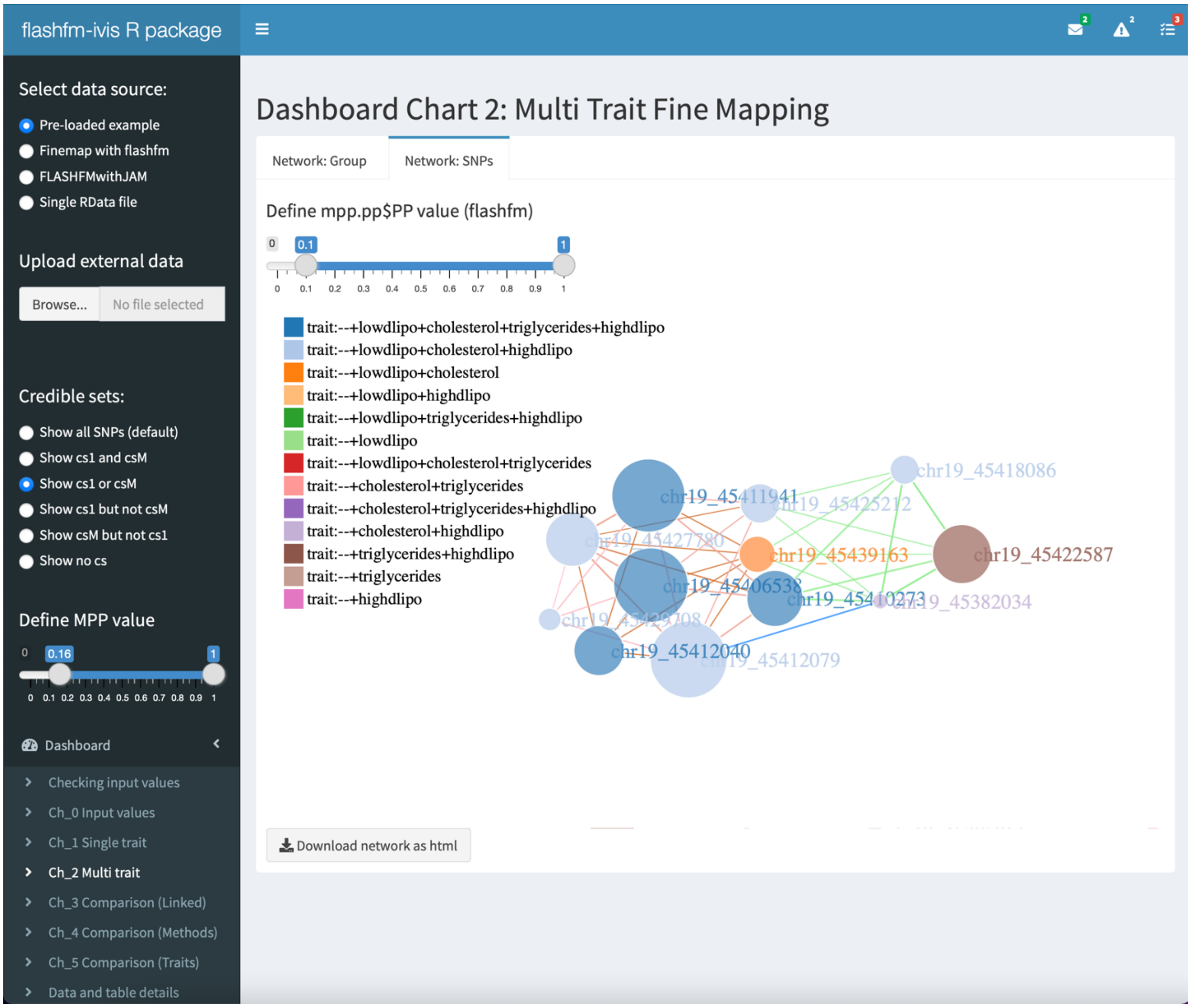

*Figure_S7: Ch_2_Multi_trait (b)*

### Ch_3 Comparison (Linked) – Multi-panel Integrated Fine-mapping regional association plots of traits for fm and flashfm

#### Coloured SNP Group– colours match those in the legend of the interactive regional association plots

a) View SNP group sizes from both single and multi-trait fine-mapping
b) The table will be adjusted automatically depending on the computer screen size, but users can also use mouse/touchpad to scroll left or right of the table.

#### Coloured and linked Manhattan plots – view and interact with both GWAS and fine-mapping results

The left panel is for single-trait fine-mapping results and the right panel is for multi-trait fine-mapping results. Each row shows individual trait results - a Manhattan plot ($-\log_{10}(p)$ against SNP position) with the following additional features:

a) Hover over a point to see SNP details (SNP ID, alleles, allele frequency, etc. Please note, if MPPg is negligible, we show it is equal to 0.) or click on "Compare data on hover" to see details for several SNPs nearby at once
b) Colour of points: SNP group membership according to fine-mapping results. SNPs belonging to the same group can be viewed as exchangeable. SNPs with MPP>0.001 are assigned to the same group if they have high LD (pairwise $r^2$>0.6) and rarely appear in a model together.
c) Size of points: proportional to fine-mapping posterior probabilities of SNP causality (referred to as MPP – Marginal Posterior Probability)
d) Click on "Box Select" or "Lasso Select" to draw a box or a lasso (free drawing of any shape) around points to focus on and fade other points. Automatically, this same set of points will become the focus in all other plots allowing simplified comparisons.
e) Click on "Zoom", then draw a box around points to zoom in and out for point selection.
f) Double click on SNP group in legend to remove all points not belonging to that group. Double click again to show all points not belonging to the SNP group
g) Click on "Pan" and then drag plot to left or right to change centre of plot.
h) Click on reset axis to res-set to default plot
i) "Download plot as a PNG" option can download the plot to the local machine
j) "Autoscale" to adjust the view of the current plot
k) Users can change their internet browser' size to adjust the view of the whole plot.

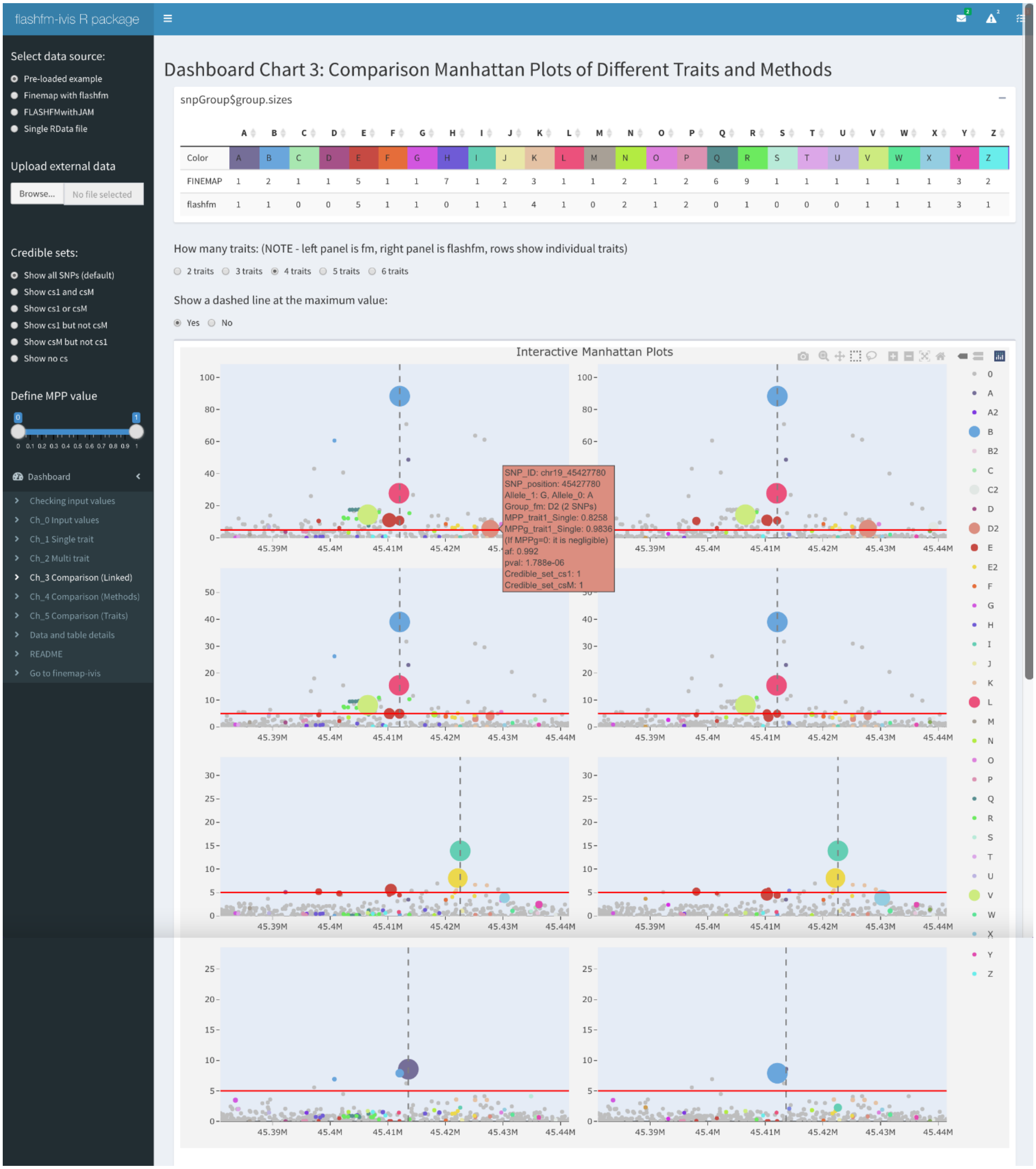


*Figure_S8: Ch_3 Comparison (linked)*

**Ch_4 Comparison (Methods) – Compare sizes and SNP overlap of credible sets**

**Radar chart of credible sets – compare sizes of credible sets between traits and methods**

a) Traits appear around the circle and the points indicate the number of SNPs in the fm credible sets (CS_1 is for 1-trait) and the flashfm credible sets (CS_M is for multi-trait)
b) Similar interactive features as other dashboards are available for users to explore the plot

**Venn diagram 1:CS_1 – overlap of fm credible sets between traits**

a) The segments show the number of SNPs that are shared between the intersecting credible sets
b) Hovering over a segment shows the SNP ids belonging to that intersection
c) The downloadable table shows the details of the credible set intersections, such as count and SNP ids. The columns can be sorted by clicking the top row options.

**Venn diagram 1:CS_M – overlap of flashfm credible sets between traits**

a) The segments show the number of SNPs that are shared between the intersecting credible sets
b) Hovering over a segment shows the SNP ids belonging to that intersection
**c)** The downloadable table shows the details of the credible set intersections, such as count and SNP ids. The columns can be sorted by clicking the top row options.

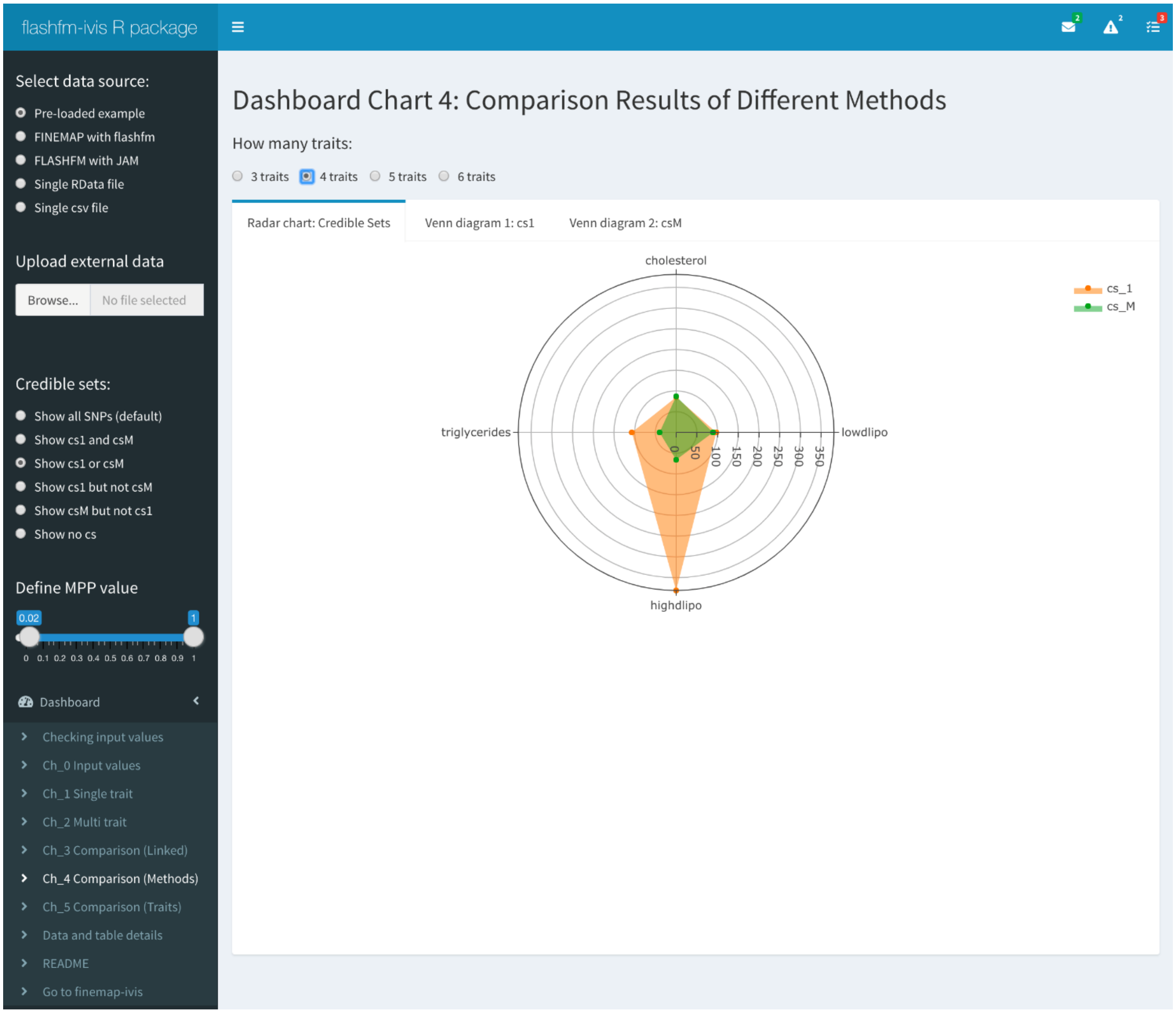

*Figure_S9: Ch_4 Comparison (Methods)(a)*

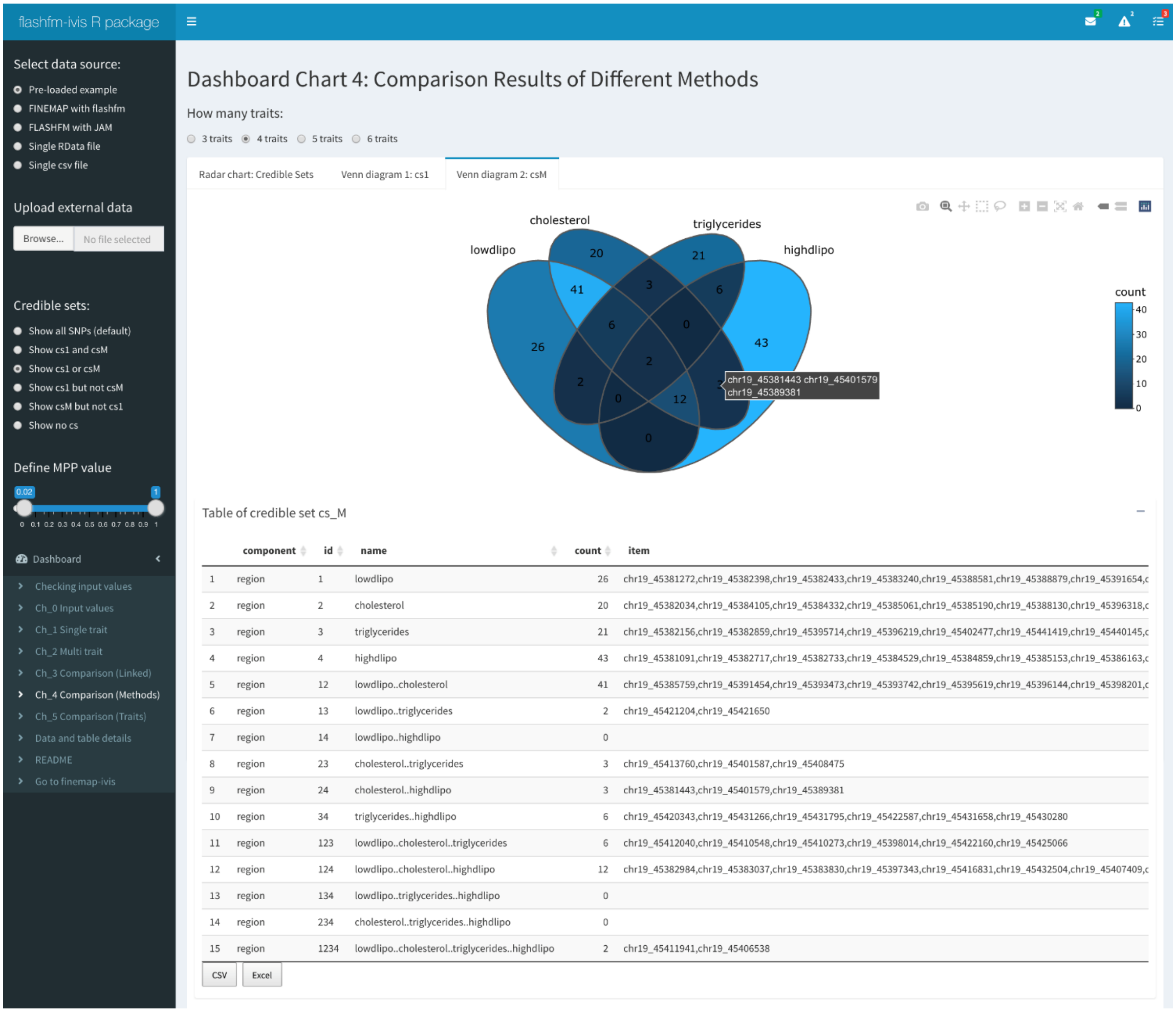


*Figure_S10: Ch_4 Comparison (Methods)(b)*

**Ch_5 Comparison (Traits) – Sankey diagrams of traits showing SNP group membership for each method**

Each Sankey diagram shows the SNPs that belong to each flashfm SNP group and to each fm SNP group. The flashfm groups tend to be subsets of the fm groups. There is an "All traits" tab and a tab for each trait. For a given fine-mapping method, SNP groups are the same for each trait. Each trait may have different SNP groups that appear in their models. To change the plot perspective (or show the names of SNPs clearly), the positions of the SNPs and groups may be moved by dragging them.

**All traits:** A combined Sankey diagram over all traits. The width of the lines joining the SNPs to their SNP group is proportional to the average MPP (over all traits, including some MPPs that are zeros or very close to zeros) for the SNP.

**Individual traits:** The width of the lines joining the SNPs to their SNP group is proportional to the trait-specific MPP for the SNP

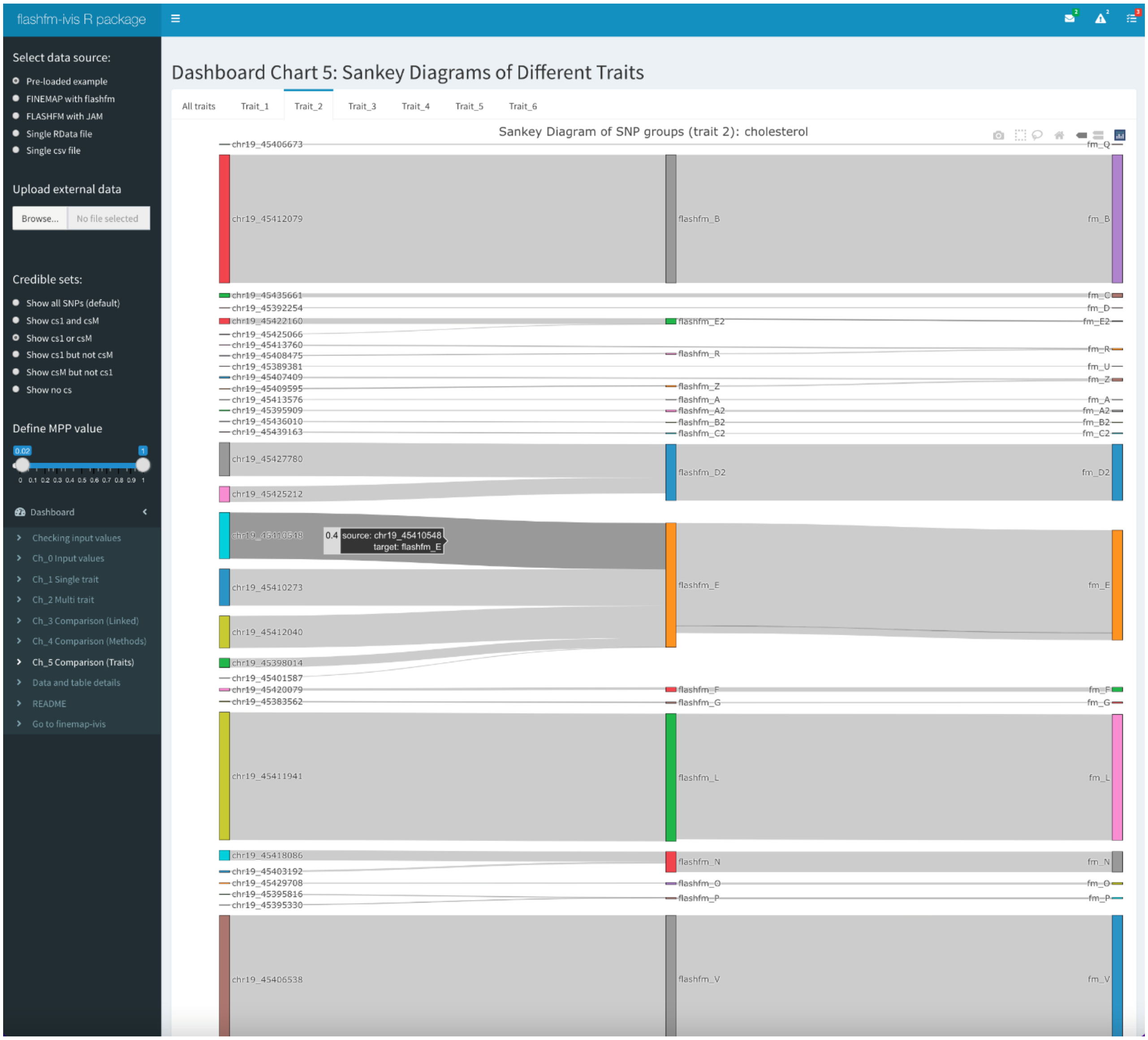


*Figure_S11: Ch_5 Comparison (traits)*

### Data and table details – shows a portion of each input object to verify input

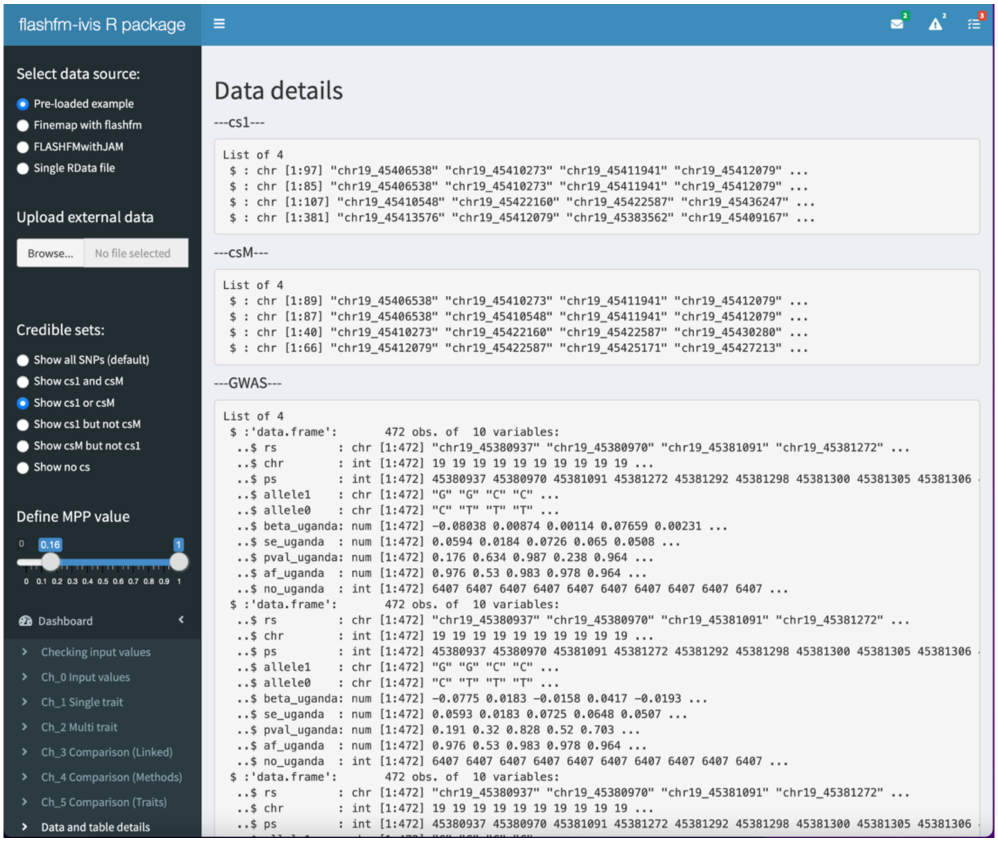

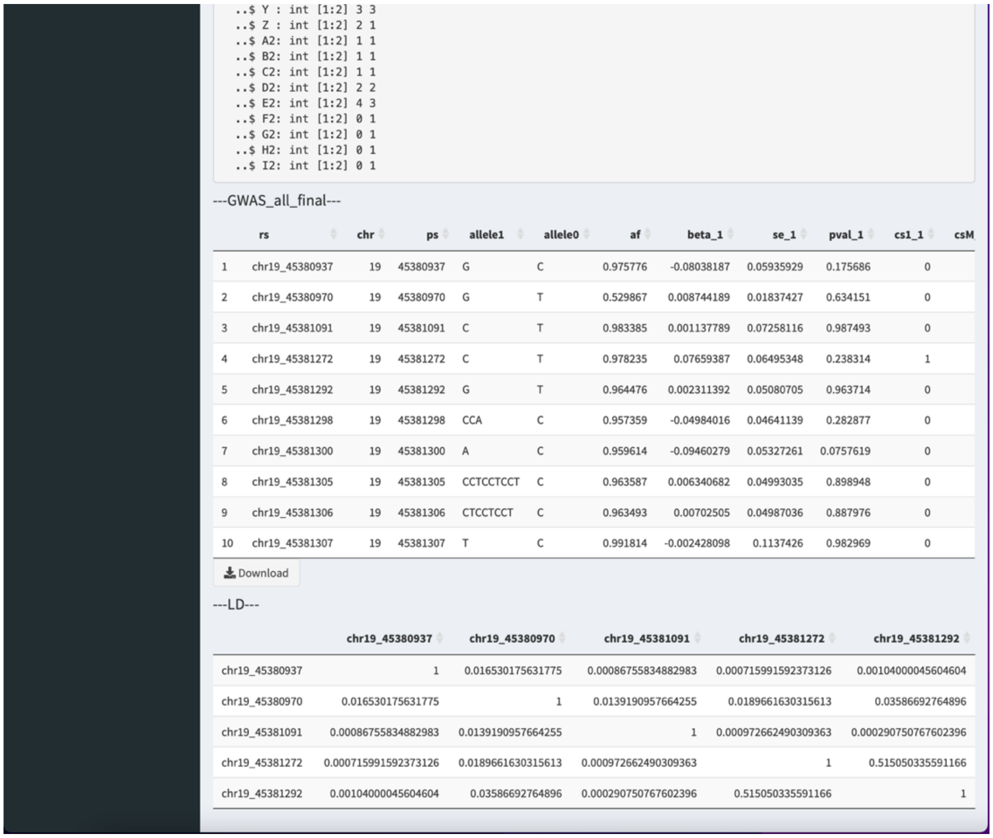

```
  ..$ Y : int [1:2] 3 3
  ..$ Z : int [1:2] 2 1
  ..$ A2: int [1:2] 1 1
  ..$ B2: int [1:2] 1 1
  ..$ C2: int [1:2] 1 1
  ..$ D2: int [1:2] 2 2
  ..$ E2: int [1:2] 4 3
  ..$ F2: int [1:2] 0 1
  ..$ G2: int [1:2] 0 1
  ..$ H2: int [1:2] 0 1
  ..$ I2: int [1:2] 0 1
```

---GWAS_all_final---

| | rs | chr | ps | allele1 | allele0 | af | beta_1 | se_1 | pval_1 | cs1_1 | csM |
|---|---|---|---|---|---|---|---|---|---|---|---|
| 1 | chr19_45380937 | 19 | 45380937 | G | C | 0.975776 | -0.08038187 | 0.05935929 | 0.175686 | 0 | |
| 2 | chr19_45380970 | 19 | 45380970 | G | T | 0.529867 | 0.008744189 | 0.01837427 | 0.634151 | 0 | |
| 3 | chr19_45381091 | 19 | 45381091 | C | T | 0.983385 | 0.001137789 | 0.07258116 | 0.987493 | 0 | |
| 4 | chr19_45381272 | 19 | 45381272 | C | T | 0.978235 | 0.07659387 | 0.06495348 | 0.238314 | 1 | |
| 5 | chr19_45381292 | 19 | 45381292 | G | T | 0.964476 | 0.002311392 | 0.05080705 | 0.963714 | 0 | |
| 6 | chr19_45381298 | 19 | 45381298 | CCA | C | 0.957359 | -0.04984016 | 0.04641139 | 0.282877 | 0 | |
| 7 | chr19_45381300 | 19 | 45381300 | A | C | 0.959614 | -0.09460279 | 0.05327261 | 0.0757619 | 0 | |
| 8 | chr19_45381305 | 19 | 45381305 | CCTCCTCCT | C | 0.963587 | 0.006340682 | 0.04993035 | 0.898948 | 0 | |
| 9 | chr19_45381306 | 19 | 45381306 | CTCCTCCT | C | 0.963493 | 0.00702505 | 0.04987036 | 0.887976 | 0 | |
| 10 | chr19_45381307 | 19 | 45381307 | T | C | 0.991814 | -0.002428098 | 0.1137426 | 0.982969 | 0 | |

Download

---LD---

| | chr19_45380937 | chr19_45380970 | chr19_45381091 | chr19_45381272 | chr19_45381292 |
|---|---|---|---|---|---|
| chr19_45380937 | 1 | 0.016530175631775 | 0.00086755834882983 | 0.000715991592373126 | 0.00104000045604604 |
| chr19_45380970 | 0.016530175631775 | 1 | 0.0139190957664255 | 0.0189661630315613 | 0.03586692764896 |
| chr19_45381091 | 0.00086755834882983 | 0.0139190957664255 | 1 | 0.000972662490309363 | 0.000290750767602396 |
| chr19_45381272 | 0.000715991592373126 | 0.0189661630315613 | 0.000972662490309363 | 1 | 0.515050335591166 |
| chr19_45381292 | 0.00104000045604604 | 0.03586692764896 | 0.000290750767602396 | 0.515050335591166 | 1 |

*Figure_S12: Data and table details*

**FINEMAP-ivis page:**

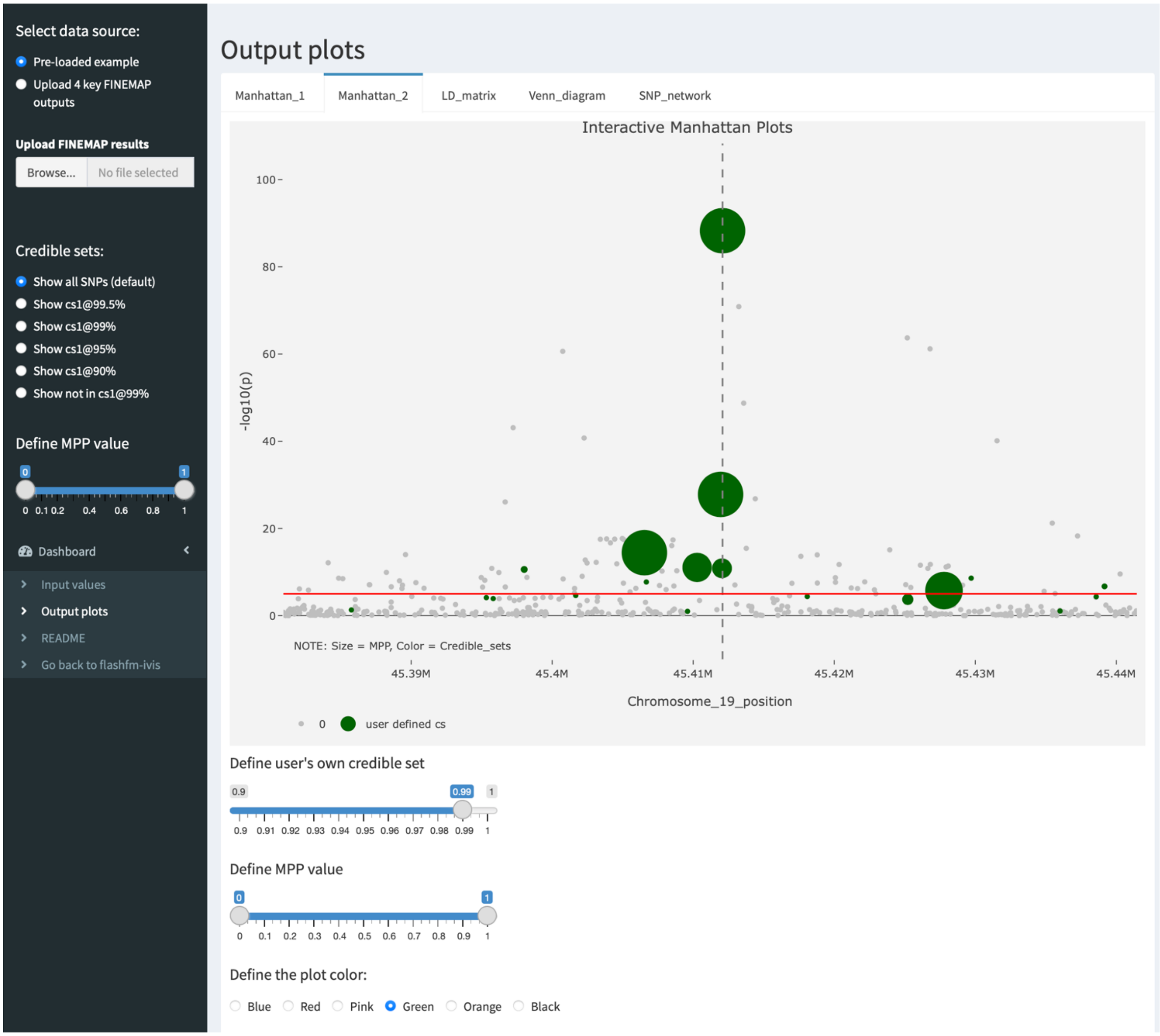


*Figure_S13: finemap-ivis*

**References (as included in Supplementary Material)**

15. Kierczak, M., Jablonska, J., Forsberg, S. K., Bianchi, M., Tengvall, K., Pettersson, M., et al. (2015). Cgmisc: enhanced genome-wide association analyses and visualization. Bioinformatics 31, 3830–3831. https://doi.org/10.1093/bioinformatics/btv426.
16. Kortemeier,E. et al. (2018) ShinyGPA: an interactive visualization toolkit for investigating pleiotropic architecture using GWAS datasets. PLoS One, 13, e0190949. https://doi.org/10.1371/journal.pone.0190949
17. Kwong, A., Boughton, A. P., Wang, M., VandeHaar, P., Boehnke, M., Abecasis, G., & Kang, H. M. (2021). FIVEx: an interactive eQTL browser across public datasets. Bioinformatics. https://doi.org/10.1093/bioinformatics/btab614.
18. Lipka, A. E., Tian, F., Wang, Q., Peiffer, J., Li, M., Bradbury, P. J., et al. (2012). GAPIT: genome association and prediction integrated tool. Bioinformatics 28, 2397–2399. https://doi.org/10.1093/bioinformatics/bts444.
19. Machiela, M. J., and Chanock, S. J. (2015). LDlink: a web-based application for exploring population-specific haplotype structure and linking correlated alleles of possible functional variants. Bioinformatics 31, 3555–3557. https://doi.org/10.1093/bioinformatics/btv402.
20. Newcombe, P. J., Conti, D. V. & Richardson, S. (2016). JAM: a scalable Bayesian framework for joint analysis of marginal SNP effects. *Genet. Epidemiol.* 40, 188–201. https://onlinelibrary.wiley.com/doi/full/10.1002/gepi.21953
21. Pruim, R. J., Welch, R. P., Sanna, S., Teslovich, T. M., Chines, P. S., Gliedt, T. P., ... & Willer, C. J. (2010). LocusZoom: regional visualization of genome-wide association scan results. Bioinformatics, 26(18), 2336-2337. https://doi.org/10.1093/bioinformatics/btq419.
22. Schilder, B. M., Humphrey, J., & Raj, T. (2021). echolocatoR: an automated end-to-end statistical and functional genomic fine-mapping pipeline. Bioinformatics. 1-4, https://doi.org/10.1093/bioinformatics/btab658.
23. Sievert, C. (2020). Interactive web-based data visualization with R, plotly, and shiny. CRC Press.
24. Verity, R., Collins, C., Card, D. C., Schaal, S. M., Wang, L., & Lotterhos, K. E. (2017). minotaur: A platform for the analysis and visualization of multivariate results from genome scans with R Shiny. Molecular ecology resources, 17(1), 33-43. https://doi.org/10.1111/1755-0998.12579
25. Wang J., Zhang Z. (2021). GAPIT Version 3: Boosting Power and Accuracy for Genomic Association and Prediction, Genomics, Proteomics & Bioinformatics, doi: https://doi.org/10.1016/j.gpb.2021.08.005.
26. Wickham, H. (2011). ggplot2. Wiley interdisciplinary reviews: computational statistics, 3(2), 180-185.
27. Ziegler,G.R. et al. (2015) Zbrowse: an interactive GWAS results browser. PeerJ Comput. Sci., 1, e3. https://doi.org/10.7287/peerj.preprints.902v1
